# Emergence and Genomic Characterization of Multidrug resistant *Candida auris* in West Africa

**DOI:** 10.1101/2022.05.24.493359

**Authors:** Rita Oladele, Jessica N. Uwanibe, Idowu B. Olawoye, Abdul-Wahab O. Ettu, Christian T. Happi

## Abstract

*Candida auris* is an emerging multidrug-resistant fungal pathogen that has become a worldwide public health threat due to limitations of treatment options, difficulty in diagnosis, and its potential for clonal transmission. Antifungal suceptibility tests and next generation sequencing were carried out on cultured isolates. Bioinformatics analysis was done using variant calling methods and genome-wide short nucleotide polymorphism (SNP) based phylogeny. Here, we report the first four cases of *C. auris* infection and colonization reported in West Africa. A total of four isolates from four reported cases of candidemia were analyzed. Three patients had fungaemia, which led to fatal invasive infection and the last patient was a likely case of colonization. Of the four patients, two had mutations which conferred resistance to the antifungal azole group and other non-synonymous mutations in hotspot genes such as ERG2, ERG11 and FKS1. Isolates from these patients clustered to clades I and IV, which indicates more than one introduction of *C*.*auris* into Nigeria. The first report of *C. auris* in Nigeria and West Africa is of public health importance as this report will aid identification, surveillance and intervention of resistant drug resistant candidiasis in the region.

## Introduction

Candida species are the predominant cause of nosocomial fungal infections, causing bloodstream infections (BSI) with significant mortality (1), therefore eliciting a major threat to intensive care unit (ICU) patients (2, 3). *Candida auris* is an emerging multidrug-resistant fungal pathogen that has become a worldwide public health threat due to limitations of treatment options, based on antifungal resistance issues and its potential for clonal transmission. In addition, difficulties in identification using conventional phenotypic and molecular techniques, the unknown population prevalence, the uncertain environmental niches, and the unclear mechanisms of spread have hindered control (4). Molecular epidemiological investigations of *C. auris* outbreaks generally show clusters of highly related isolates, supporting local and ongoing transmission. The analysis of outbreaks and individual cases has also revealed genetic complexity, with isolates from different clades detected in Germany, United Kingdom, and United States, suggesting multiple introductions into these countries, followed by local transmission (5). Since it was first isolated from an ear canal sample of a Japanese patient in 2009, it has been isolated in several countries on five continents across the world. Mortality rates have varied significantly among geographic regions from Asia, far East and the United states have reported a 50% mortality rate this is in contrast to Venezuela where survival rate is 72% (6). In Nigeria, the prevalence of *C. auris* is unknown because no cases of colonization or infection have been reported (6). Yet given its geographical location, with frequent travel of the population for business and medical purposes to India, South Africa, USA and UK to the west, the country is at high risk for the spread of this pathogen.

In this study we report the first four cases of *C. auris* infection and colonization in Nigeria. This is also the first report of *C. auris* outbreak in West Africa, to the best of our knowledge.

### Case reports

Four cases of invasive candidiasis were reported from four different facilities in Nigeria: two from ICUs in two private facilities in Lagos Nigeria; one from a tertiary hospital in Ibadan (which is approx 200 km from Lagos) and the last, a female from a tertiary hospital in Lagos. Candida spp was isolated from blood culture in all four patients. Clinical specimens were plated on Sabouraud dextrose agar plates. The isolates were initially identified phenotypically as non-*Candida albicans*. But, with deteriorating clinical conditions of the patients, the isolates were sent to the Lagos University Teaching Hospital (LUTH) which had vitek-2 (9.01 version) and yeast cards routinely available for identification and antifungal susceptibility testing. The organisms were identified as *C. auris*. Table 1 below is a summary patients demographics, clinical presentation, diagnosis and outcome of the cases.

**Table 1:**
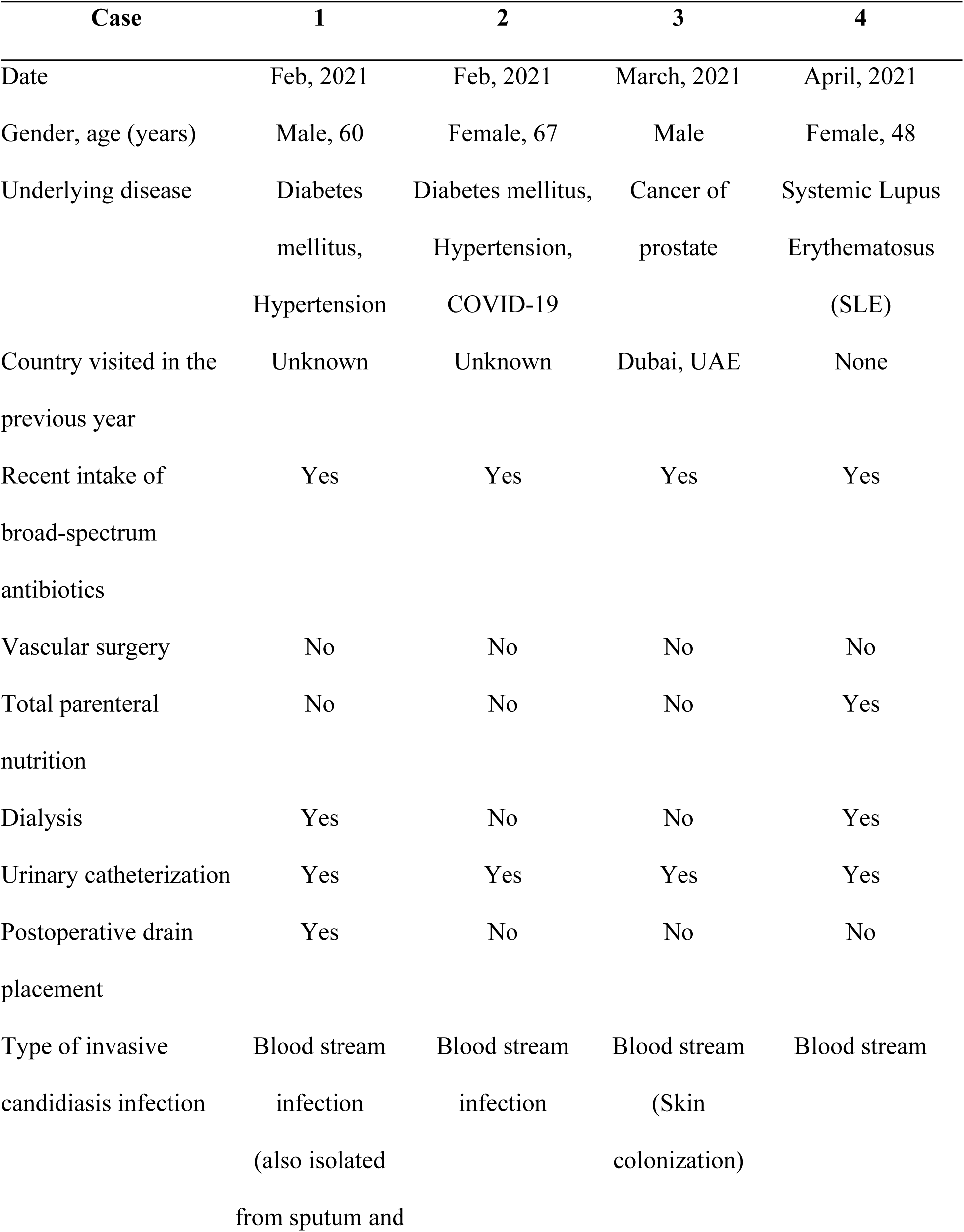
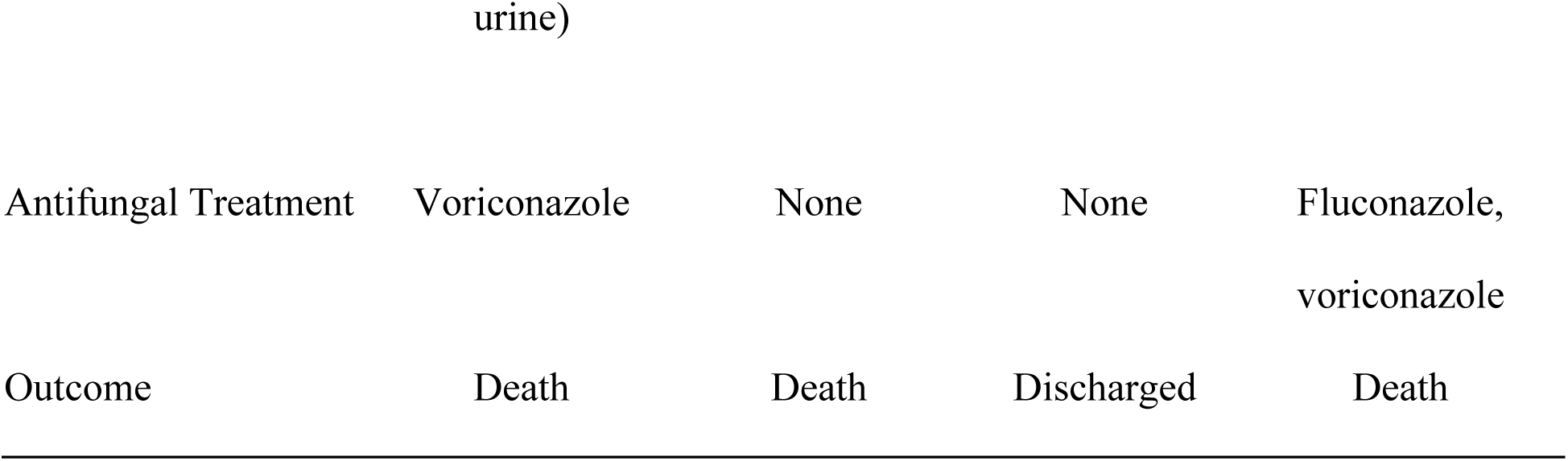
Demographics and clinical characteristics of reported cases

### Antifungal susceptibility test (AFST)

To examine resistance levels to the available drugs, antifungal susceptibility tests were performed using: Fluconazole, Voriconazole, Posaconazole, Amphotericin B, Caspofungin, Micafungin, and Anidulafungin. Results exhibited high minimum inhibitory concentrations (MIC) to triazoles and amphotericin B in cases 1, 2 and 3 (Table 2).

**Table 2:** Antifungal susceptibility test results of *Candida auris* isolates

| **Antifungal (minimal inhibitory concentration mg/L)** |  |  |  |  |  |  |  |
| --- | --- | --- | --- | --- | --- | --- | --- |
|  | **Fluconazole** | **Voriconazole** | **Posaconazole** | **Amphotericin B** | **Caspofungi** | **Micafungir** | **Anidulafung** |
|  |  |  |  |  |  |  | **n** |
| Tentative<br>MIC<br>breakpoints* | ≥32 | N/A** | N/A** | ≥2 | ≥2 | ≥4 | ≥4 |
| Case 1 | 16 | 0.06 | - | 1 | 0.06 | 0.06 | 0.12 |
| Case 2 | ≥32 | 1 | - | 0.5 | 0.06 | 0.06 | 0.12 |
| Case 3 | 1 | 0.02 | - | 0.5 | 0.06 | 0.06 | 0.12 |
| Case 4 | NA | NA | NA | NA | NA | NA | NA |

\*Breakpoints are defined based on CDC guidance for *C. auris* MIC interpretation (7)
\*\*Fluconazole susceptibility is used as a surrogate for second generation triazole susceptibility assessment
NA - Not available

## Methods

### DNA extraction and library preparation

All isolates were subcultured for DNA extraction and WGS. DNA was extracted using the Plant/Fungi DNA isolation kit (Norgen Biotek CORP, Canada). DNA samples were quantified using Qubit fluorometer (ThermoFisher Scientific) using dsDNA High sensitivity assay. Sequencing libraries were prepared using the Nextera DNA flex preparation kit (Illumina, USA) adopted from a previous study (8).

### Whole genome sequencing and bioinformatics analysis

Four isolates were sequenced on the Illumina MiSeq using Illumina DNA flex library preparation kit and the v2 300 cycle kit. FASTQ files that were generated from the MiSeq underwent quality control checks using FASTQC and further improved with fastp (9). The resulting FASTQ paired-end files were mapped against the *Candida auris* reference genome (GCF_002775015.1) which is 12.7 millionbase pairs (Mbp) long, and contains seven (7) chromosomes using BWA MEM (10) algorithm. Duplicate reads were marked and removed from the resulting mapped reads using SAMtools and a coverage plot was generated by normalizing the maximum read depth at 14,964 base pair window (15 kbp).

Variants were called using Freebayes (11) v1.3.2 by using variant filtering parameters of MQM (Mean mapping quality) > 45, DP (Read depth) > 10. The VCF file was annotated with snpEff (12) v 5.0e and filtered using SnpSift (13) v5.0e to focus on specific genes of interest *ERG11, ERG3*, ERG2, ERG6, *FKS1*, FKS2 and FUR1 with non synonymous mutations excluding mutations in upstream, downstream and intergenic regions.

### Phylogenetic analysis

We analyzed 37 *C. auris* whole genomes previously studied (5), from which have already been classified into four (4) distinct lineages, in addition with the four (4) isolates from this study. The VCFs generated from the freebayes were merged with BCFtools (14) v1.10.2 taking only SNPs into account. Genome-wide SNP phylogenetic analysis was performed on all samples taking into account unambiguous SNPs (n=174,758) with an ultrafast bootstrap value of 1000 and a generalized time reversible model using IQ-TREE (15) which was visualized on figtree.

## Results

The average coverage of *C. auris* reads that mapped against the reference genome after quality improvement with fastp were 44.15x, 51.85x, 59.23x and 67.26x respectively with all seven (7) chromosomes sequenced as seen in Figure 1.

**Figure 1.**
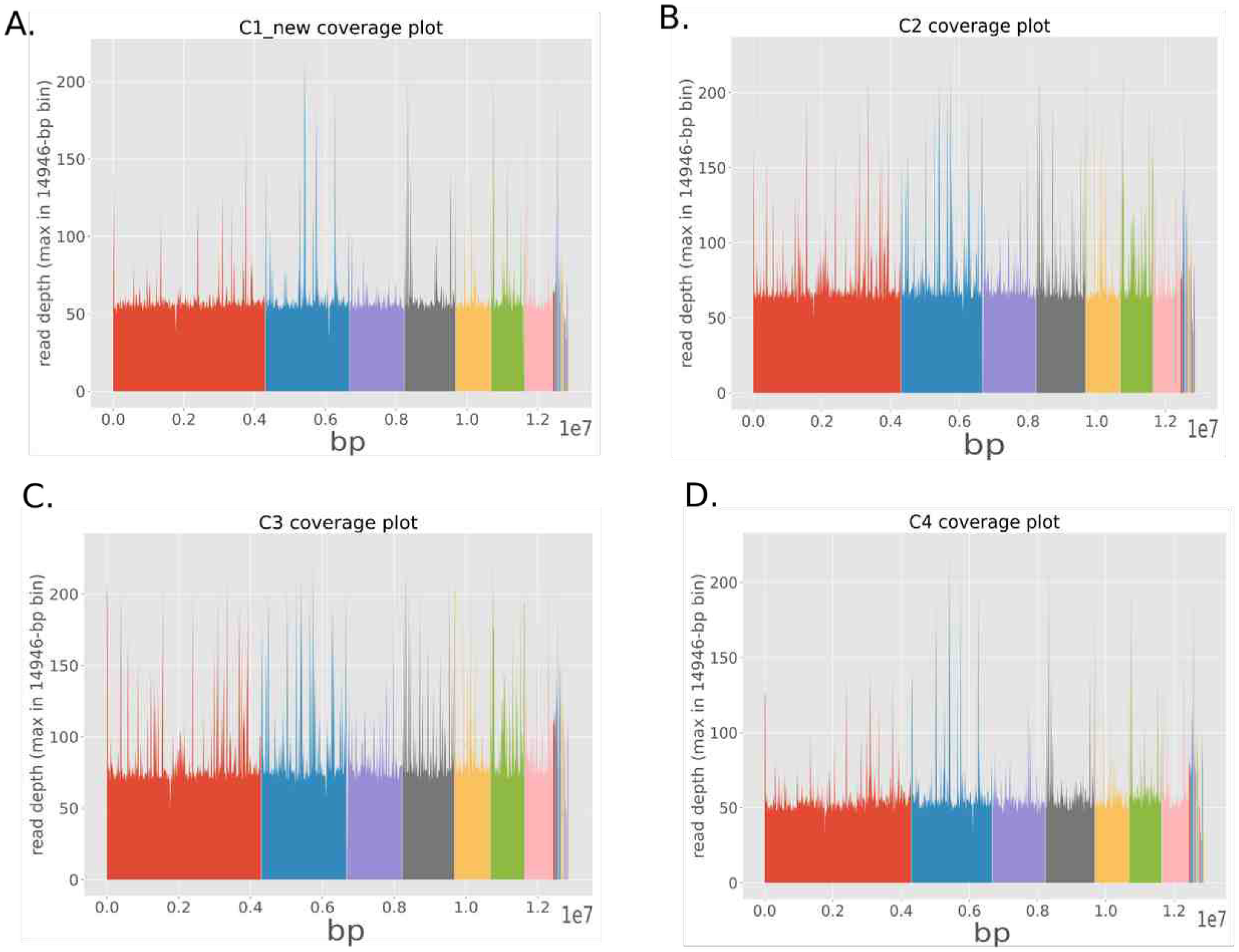
Coverage plot of whole genome sequencing of *C. auris* isolates from case one (A), case two (B), case three (C) and case four (D) patients. Sequencing depth was calculated by plotting the maximum average per 15 kilobase pair (kbp) window across the genome size, which is the x-axis raised to the power of 7. Each colour represents the different chromosome from 1 to 7 (left to right).

### Variant analysis

Following genome characterization and assembly, specific genotypes conferring antifungal resistance were investigated in hotspot genes such as *ERG11, ERG3*, ERG2, ERG6, *FKS1*, FKS2 and FUR1. All isolates had non-synonymous mutations (N335S, E343D, Y132F, L125F, and K177R) in the *ERG11* gene. Case 2 had additional non-synonymous mutations (L148I, R937S, I701V and I694V) in the *FKS1* gene, and case 4 with another non-synonymous mutation (E39D) in the *ERG2 gene* (Table 3). Cases 1 and 4 patients were both placed on Fluconazole, with voriconazole as an added medication for the latter, however all patients except case 3 died during treatment.

**Table 3:**
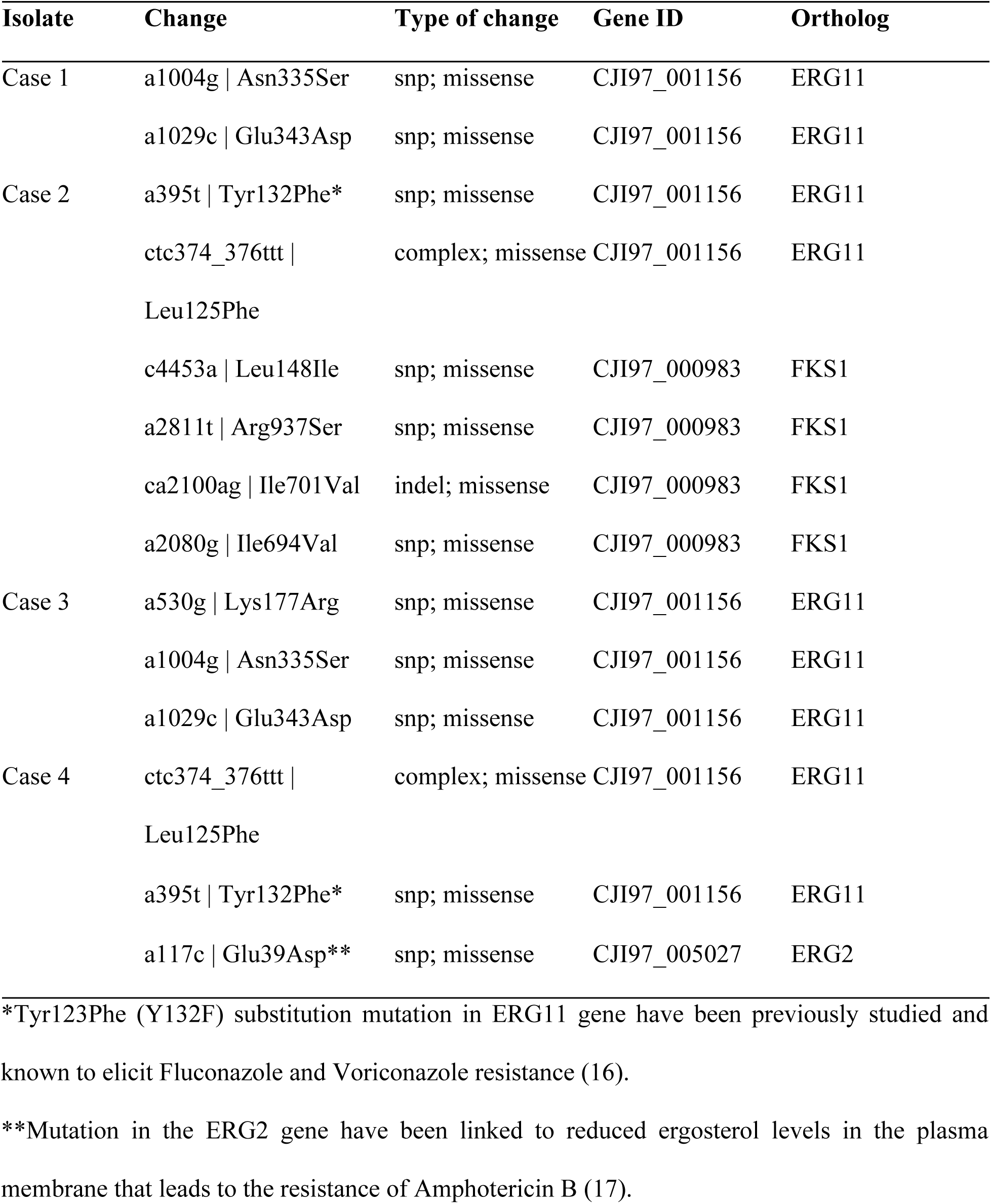
Detected polymorphisms in hotspot genes that are associated with drug resistance in *C. auris* in the four (4) cases/isolates

### Phylogenetic analysis

From phylogenetic analysis, cases 2 and 4 which had confirmed mutations (*ERG11:*Y132F) conferring drug resistance to azoles clustered with clade 1, whilst cases 1 and 3 clustered with clade 4 (Figure 2).

**Figure 2.**
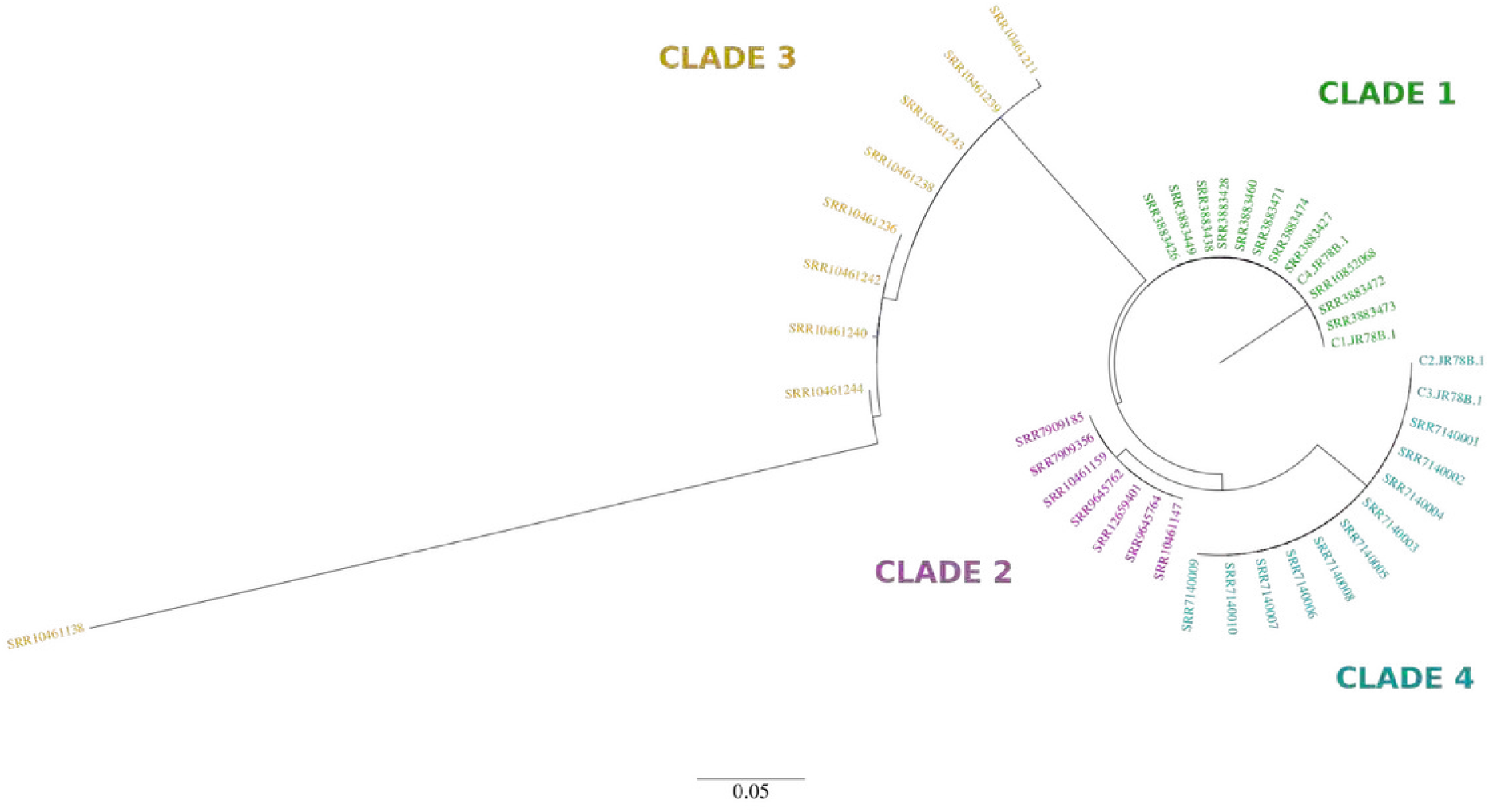
Maximum-likelihood phylogeny of 41 *C. auris* genomes (including 4 from this study) showing four distinct phylogeographical clades from samples all around the world. Genomes from this study cluster with clades 1 and 4.

Clade 1 is the most globally spread of *C. auris* which have been found in 11 countries such as Canada, France, Germany, India, Kenya, Pakistan, Saudi Arabia, United Kingdom, UAE and the USA. However, clade 4 is mostly found in the Americas and it has been discovered in Colombia, Israel, Panama, USA and Venezuela (5).

## Discussion

In this study, we report the first cases of *C. auris* in Nigeria which also happens to be the first reported cases in the West African region. Antifungal susceptibility tests in addition to whole genome sequence analysis revealed high resistance levels to triazoles and amphotericin B which were supported by non-synonymous mutations in *ERG2, ERG11* and *FKS1* genes.

*C. auris* causes outbreaks in hospitals, and cases have been reported in Colombia, Venezuela, Israel, South Africa, and the United Kingdom (18 - 22). Within weeks of an index patient joining the facility, patients in a long-term care facility can get colonized with *C. auris* (21). It has been identified among patients exposed to the ventilator and long term acute care hospitals, and likewise among those who have received healthcare in countries with increasing *C. auris* transmission, immunocompromised, recent broad-spectrum antibiotic or antifungal and/or central venous catheter (23, 24). It can be misidentified for *Candida haemulonii, Candida famata, Saccharomyces cerevisiae*, and *Rhodotorula glutinis* with the analytical profile index strips or the VITEK (21, 23).

As the part of this study, 600 Candida isolates from different sites in Nigeria in 2019, were sent for sequencing in search of possible *C. auris*, 210 were successfully sequenced and none was *C. auris*. In discussion with collaborators at CDC Atlanta, information of a case of *C. auris* isolated from a Nigerian child who was in New York for Medical care flagged off alarm of the possibility of the infection being acquired from Nigeria. This was at the peak of the first wave of COVID-19, thus investigating it was not feasible there. Here, we report four recent cases of *C. auris* observed in Nigeria over a three-month period. Previous genomic studies have revealed that the four (4) major distinct clades of *C. auris* emerged independently from different geographical points across the world (25), and our phylogenetic analysis infer that there might have been at least two different introductions of *C. auris* in Nigeria that emerged independently of each other. In addition, we did not find known mutations in hotspot genes which confer resistance to Echinocandins such as Caspofungin, Micafungin and Anidulafungin. The genetic/molecular findings correlate with the Antifungal susceptibility tests done earlier.

As *C. auris* is a multidrug-resistant pathogen and is prone to misidentification by available conventional methods; the true burden of this problem in Nigeria remain unknown, given that most routine clinical laboratory in the country limits *Candida sp* identification to just *C. albicans* and non-*C*.*albicans*. This is further compounded by the poor availability and accessibility of antifungals in the country (only fluconazole, itraconazole, voriconazole and AmphotericinB deoxycholate are licensed in Nigeria), as three out four of the patients diagnosed, succumbed to their illnesses and passed away. This calls for rapid and accurate molecular diagnosis of *C. auris* in hospitals as the genomes generated in this study can be instrumental in developing one.

## Conclusion

In this study, we report the first evidence of *C. auris* infections in Nigeria and West Africa, which were further confirmed using whole genome sequencing. All of these isolates were seen to contain substitution mutations in hotspot genes responsible for antifungal resistance to triazoles and amphotericin B which are of serious global health concern, especially in hospitals. This work also shows how healthcare systems in West Africa can hugely benefit from hospital and academic research collaboration in identifying and diagnosing pathogens that poses a huge threat to human lives.

## Data Availability

The FASTQ files of isolates generated from this study are availabe in the NCBI Sequence read archive (SRA) under BioProject accession number PRJNA838244.

## Notes

## Acknowledgements

We would like to thank Meghan M. Lyman and Tom Chiller from the Mycotic Diseases branch, Centres for Disease Control (CDC), USA for their collaboration and continued support. We also thank the Lagos State Hospital Management Board.

## Disclaimer

The funder had no role in the design and conduct of the study; collection, management, analysis, and interpretation of the data; preparation, review, or approval of the manuscript; and decision to submit the manuscript for publication.

## Funding

This work is made possible in ACEGID, Redeemer’s University, Ede, through support from a cohort of generous donors through TED’s Audacious Project, including the ELMA Foundation, MacKenzie Scott, the Skoll Foundation, and Open Philanthropy. This work was also supported by grants from the National Institute of Allergy and Infectious Diseases (https://www.niaid.nih.gov), NIH-H3Africa (https://h3africa.org) (U01HG007480 and U54HG007480), the World Bank (ACE-IMPACT project), the Rockefeller Foundation (Grant #2021 HTH), the Africa CDC through the African Society of Laboratory Medicine (ASLM) (Grant #INV018978), and the Science for Africa Foundation.

## Conflict of interest

The authors: No reported conflicts of interest.

